# Low-affinity CAR T cells exhibit reduced trogocytosis, preventing fratricide and antigen-negative tumor escape while preserving anti-tumor activity

**DOI:** 10.1101/2021.12.05.471117

**Authors:** Michael L. Olson, Erica R. Vander Mause, Sabarinath V. Radhakrishnan, Joshua D. Brody, Aaron P. Rapoport, Alana L. Welm, Djordje Atanackovic, Tim Luetkens

## Abstract

Chimeric antigen receptor (CAR) T cells using the high-affinity CD19 binding domain FMC63 are an effective treatment for patients with relapsed and aggressive B cell lymphoma. However, antigen loss and poor CAR T cell persistence remain common causes for relapse in these patients. Using primary patient samples, we now show that FMC63-based CAR T cells confer rapid antigen loss in all major tumor types currently approved for treatment with CD19 CAR T cells via trogocytosis, the stripping of antigen from tumor cells by CAR T cells. We show that CAR T cell-mediated trogocytosis can be dramatically reduced across a wide range of B cell malignancies by replacing FMC63 with a low affinity CD19 antibody. This reduction in trogocytosis does not alter the direct anti-tumor activity of CD19 CAR T cells but prevents the emergence of antigen-negative tumor cells and significantly increases CAR T cell viability by reducing fratricide of CD19 CAR T cells following trogocytosis.

**TEASER:** A reduction in CAR affinity does not affect tumor killing but prolongs T cell persistence and prevents antigen-negative tumor escape.

## INTRODUCTION

Chimeric antigen receptor (CAR) T cells, specifically CAR T cells targeting CD19, have greatly improved outcomes for patients with B cell lymphoma; however, disease relapse remains a common occurrence in many patients (*1-4*). Different mechanisms of relapse have been demonstrated, such as CD19 open-reading frame mutations, upregulation of coinhibitory ligands, and CAR T cell exhaustion (*5-13*). In many relapsed patients no clear immune escape mechanisms could be identified and additional processes are likely to play an important role.

Trogocytosis is a process originally described in the TCR context whereby TCR internalization upon binding of peptide-MHC results in stripping of MHC from antigen presenting cells and subsequent expression on the surface of the recipient T cell (*14*). Trogocytosis has since been observed in many facets of natural immunity and plays an important role in immune cell development and function (*15-22*). Recently, CAR T cells were also shown to exhibit trogocytosis when co-cultured with different types of cancer cells, including CAR T cells based on the high-affinity CD19 binding domain FMC63, which were shown to strip CD19 from lymphoma cells and incorporate it into their own plasma membrane (*23*). In addition, it was shown that CAR T cells engineered to express CD19 can be killed by CD19 CAR T cells suggesting that such fratricide might also occur spontaneously after CAR T cells acquire CD19 by trogocytosis (*23*). Importantly, to date the actual effect of trogocytosis on fratricide remains unknown and no strategies exist to limit trogocytosis in the CAR T cell context to prevent the emergence of antigen-negative tumor cells.

## RESULTS

### CLL cells become CD19-negative in vitro and in vivo after CD19 CAR T cell treatment

To quantify trogocytosis-induced CD19 loss across B cell malignancies, we first generated CAR T cells targeting CD19 based on the high-affinity scFv FMC63 used in all three FDA-approved CD19 CAR T cell products, as well as non-targeting CAR T cells that lack an a binding domain (ΔscFv; Fig. 1A). After culturing high-affinity CD19 CAR T cells with primary chronic lymphocytic leukemia (CLL) cells for 4 hours, CD19 had become undetectable on the surface of nearly all viable CLL cells (Suppl. Fig. 1A-B). Intracellular staining (Suppl. Fig. 1B) and western blot analysis (Suppl. Fig. 1C) revealed a loss of total CD19 protein from CLL cells following coculture, suggesting that loss of CD19 surface expression was not due to internalization by the CLL cells. We next utilized a short-term, dose limited ALL mouse model (Fig. 1B) with incomplete tumor elimination (Fig. 1C) in order to quantify the effect of CD19 CAR T cell-mediated trogocytosis on established tumors. Flow cytometry analysis of lymph nodes revealed complete CD19 loss on the surface of most remaining live NALM6 cells treated with CD19 CAR T cells, when compared to those treated with ΔscFv CAR T cells (Fig. 1D). The observed effect was comparable to in vitro co-culture assays, confirming that CAR T cell-mediated trogocytosis robustly confers antigen-loss in B-ALL cell.

**Figure 1.**
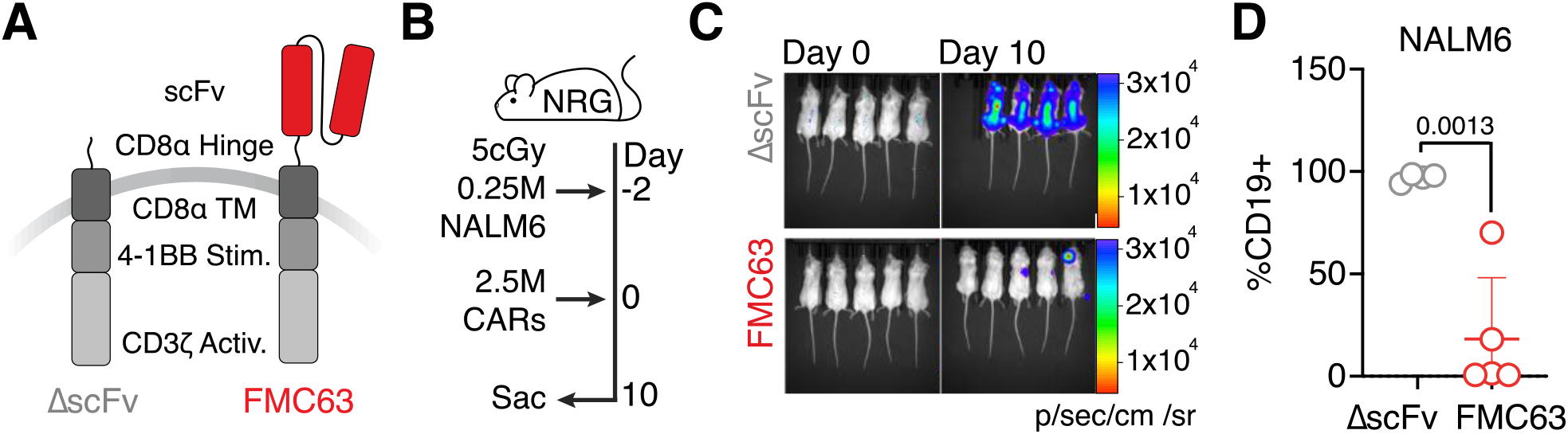
CD19 CAR T cells strip CD19 from B cell lymphoma cells *in vivo*. **(A)** Schema of ΔscFv and CD19 targeted CARs. **(B)** Schema of trogocytosis mouse model. **(C)** IVIS signal in NALM6-luc bearing mice treated with ΔscFv (top) or CD19 (bottom) CAR T cells. **(D)** %CD19+ NALM6-luc cells in the bone marrow of ΔscFv (grey) or CD19 (red) CAR T cell treated NSG mice. Data represent mean ± SD from ΔscFv (N=4) treated and FMC63 (N=5) treated mice. Statistical significance was determined by two-sided Student’s *t* test.

### CD19 is transferred rapidly from CLL cells to CAR T cells

The mechanism causing trogocytosis and its kinetics remain poorly understood, complicating the clinical investigation of trogocytosis as a potential driver of persistent disease and reduced CAR T cell persistence. We therefore next determined the kinetics of CD19 CAR T cell-mediated trogocytosis and found that a small CD19-negative population of CLL cells becomes detectable only 30 minutes into a coculture with CD19 CAR T cells and grew to include >90% of CLL cells by 5 hours of coculture (Fig. 2A). We also observed the appearance of CD19^+^ CD19 CAR T cells as early as 5 minutes into coculture with CLL cells. This population remained at this level for the first 30 minutes and then started to diminish over the next 4.5 hours, potentially due to fratricide (Fig. 2B). These data indicate that trogocytosis is a very rapid process, likely resulting from few cellular interactions and potentially resulting in the persistent targeting of CD19 CAR T cells. We hypothesize that the rapid transfer of antigen and its immediate effect on CAR T cells will likely be difficult to capture in a clinical setting in the absence of a more comprehensive mechanistic understanding, e.g. to allow its targeted inhibition.

**Figure 2.**
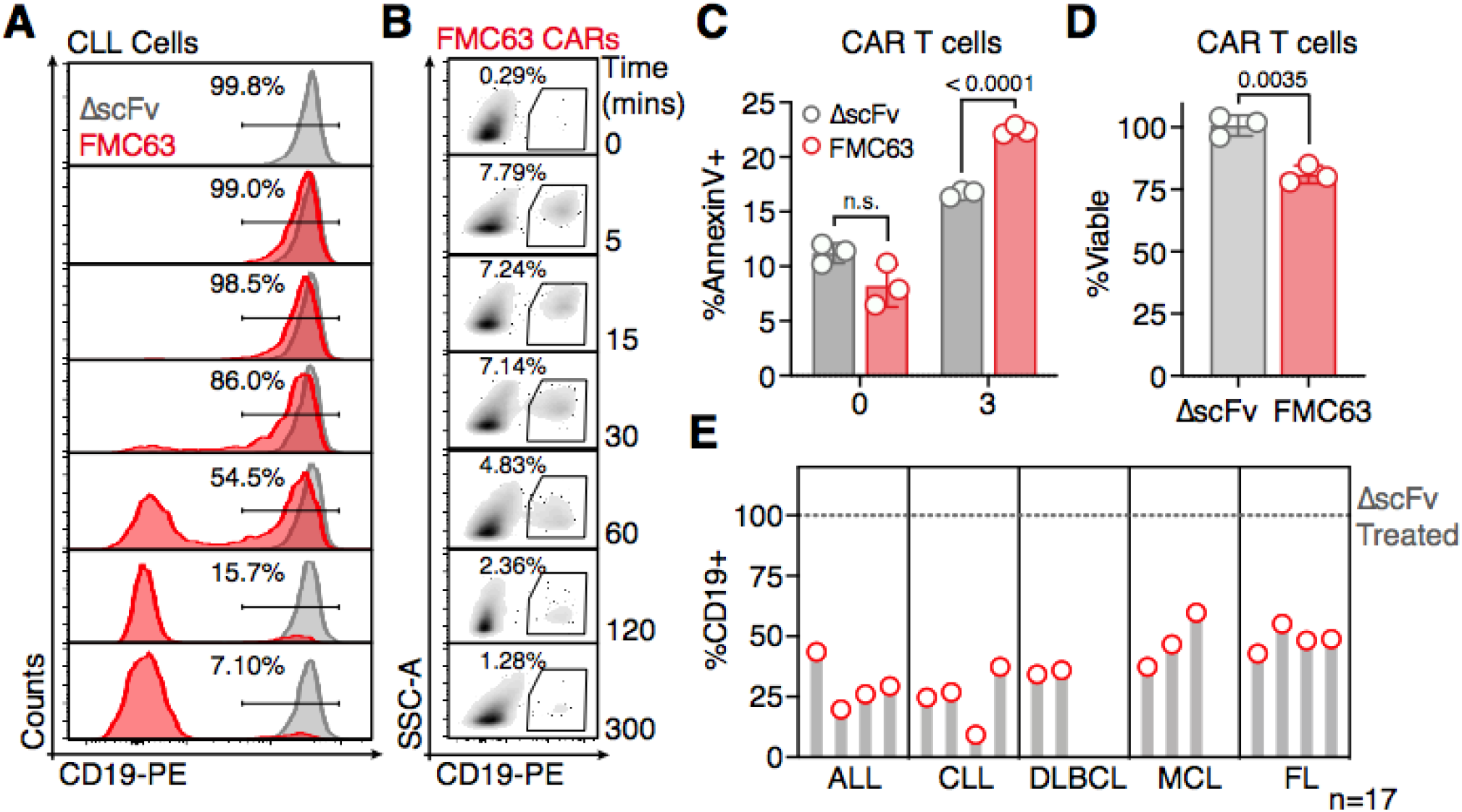
CD19 CAR T cells rapidly strip CD19 from primary B cell lymphomas, resulting in fratricide. **(A)** CD19 surface expression on primary CLL cells following coculture with ΔscFv (grey) or CD19 (red) CAR T cells at different timepoints. **(B)** CD19 surface expression on CD19 CAR T cells following coculture with primary CLL cells at different timepoints. **(C)** AnnexinV staining of ΔscFv (grey) or CD19 (red) CAR T cells following a 0- or 3-hour coculture with primary CLL cells at an effector:target ratio of 4:1. Data represent mean ± SD from three replicates. Statistical significance was determined by two-sided Student’s *t* test. **(D)** Viability of ΔscFv (grey) and CD19 (red) CAR T cells following a 4-hour coculture with primary CLL cells at an effector:target ratio of 4:1. Data represent mean ± SD from three replicates. Statistical significance was determined by two-sided Student’s *t* test. **(E)** CD19 expression on primary ALL, CLL, DLBCL, MCL and FL cells following a 4-hour coculture with CD19 CAR T cells at an effector:target ratio of 4:1. Values are normalized to CD19 expression following coculture with ΔscFv CAR T cells.

### Trogocytosis results in reduced CAR T cell viability

Previous work argued that CAR T cell-mediated trogocytosis may result in poor CAR T cell persistence due to fratricide of CD19^+^ CD19 CAR T cells. However, this was not demonstrated to occur as a direct result of coculture with CD19^+^ lymphoma cells but following transduction of CAR T cells with a constitutive CD19 expression construct (*23*). To quantify targeting of CD19 CAR T cells following trogocytosis by other CD19 CAR T cells, we assessed the binding of the early apoptosis marker Annexin V to ΔscFv and CD19 CAR T cells following coculture with primary CLL cells. CD19 CAR T cells exhibited significantly greater levels of Annexin V staining when compared to ΔscFv CAR T cells upon coculture with CLL cells (Fig. 2C) and the total viability of CD19 CAR T cells was significantly reduced after a 4 hr coculture with primary CLL cells (Fig. 2D).

### Major B cell malignancies are susceptible to CAR T cell-mediated trogocytosis

Considering the substantial effect of trogocytosis on the emergence of antigen-negative tumor cells as well as CAR T cell viability, we next sought to determine the relative susceptibility to trogocytosis of different B cell malignancies approved or under clinical investigation for treatment with CD19 CAR T cell products using the high-affinity FMC63 binding domain. We analyzed CD19 expression on a comprehensive set of primary ALL, CLL, diffuse large B cell lymphoma (DLBCL), mantle cell lymphoma (MC) and follicular lymphoma (FL) cells (N=17) following coculture with CD19 CAR T cells. While we did observe some differences in the level of trogocytosis between tumor types, all B cell malignancies were susceptible to trogocytosis when cocultured with CD19 CAR T cells (Fig. 2E), indicating that trogocytosis is a universal process and not dependent on target cell biology.

### Expansion, activation, and anti-tumor activity of CD19 CAR T cells are independent of affinity

Trogocytosis has previously been shown to correlate with Fc receptor-IgG affinity and TCR-peptide-MHC avidity(*24, 25*). We therefore hypothesized that CAR affinity may have a similar relationship to CAR T cell-mediated trogocytosis. While optimal CAR affinity for maximal cytotoxic function remains an active area of research, it has been shown that the cytotoxic activity of CARs can be maintained over a wide range of affinities depending on the target antigen (*26-28*). Binding domains currently in clinical use, such as FMC63, likely exceed the threshold affinity for maximal cytotoxic activity by several orders of magnitude. We therefore hypothesized that it may be possible to separate therapeutic CAR T cell function from trogocytic activity by modulating CAR binding domain affinity. A low-affinity CAR targeting CD19 (“CAT”) with ∼50-fold lower affinity than the clinically approved FMC63-based CAR was recently shown to exhibit enhanced efficacy and persistence in a mouse model as well as two clinical trials (*29, 30*). However, the mechanism behind this enhanced persistence and anti-tumor activity remains unknown. We generated non-binding (ΔscFv), low affinity (CAT) and high affinity (FMC63) CD19 CAR T cells in order to determine the impact of CAR affinity on trogocytosis (Fig. 3A). CAT and FMC63 CAR T cells both exhibited similar expansion during CAR T cell production (Suppl. Fig. 2A). Importantly, CAT and FMC63 were expressed at similar levels on the CAR T cell surface, suggesting that any difference in CAR function between CAT and FMC63 CAR T cells would be unrelated to CAR expression (Suppl. Fig. 2B-C). FMC63 CAR T cells bound greater amounts of recombinant CD19 than CAT CAR T cells, likely due to their higher affinity (Fig. 3B). To separate effects that may arise from differences in affinity rather than differences in trogocytosis, we next determined whether CAR affinity impacts CAR T cell activation and expansion when stimulated by antigen without allowing for trogocytosis by using CD19-coated beads instead of target cells. We found that stimulation of FMC63 and CAT CAR T cells with CD19-coated beads resulted in equal levels of IFN*γ* production and CAR T cell expansion when compared to unstimulated controls, suggesting that CAR ligation results in similar short-term cellular outcomes, regardless of CAR affinity (Fig. 3C-D). Finally, in a short-term cytotoxicity assay, we found that FMC63 and CAT CAR T cells both effectively lysed primary CLL cells without a significant difference in cytotoxicity by the two CARs (Fig. 3E), indicating that the reduction in affinity did not lead to significantly altered CAR T cell function.

**Figure 3.**
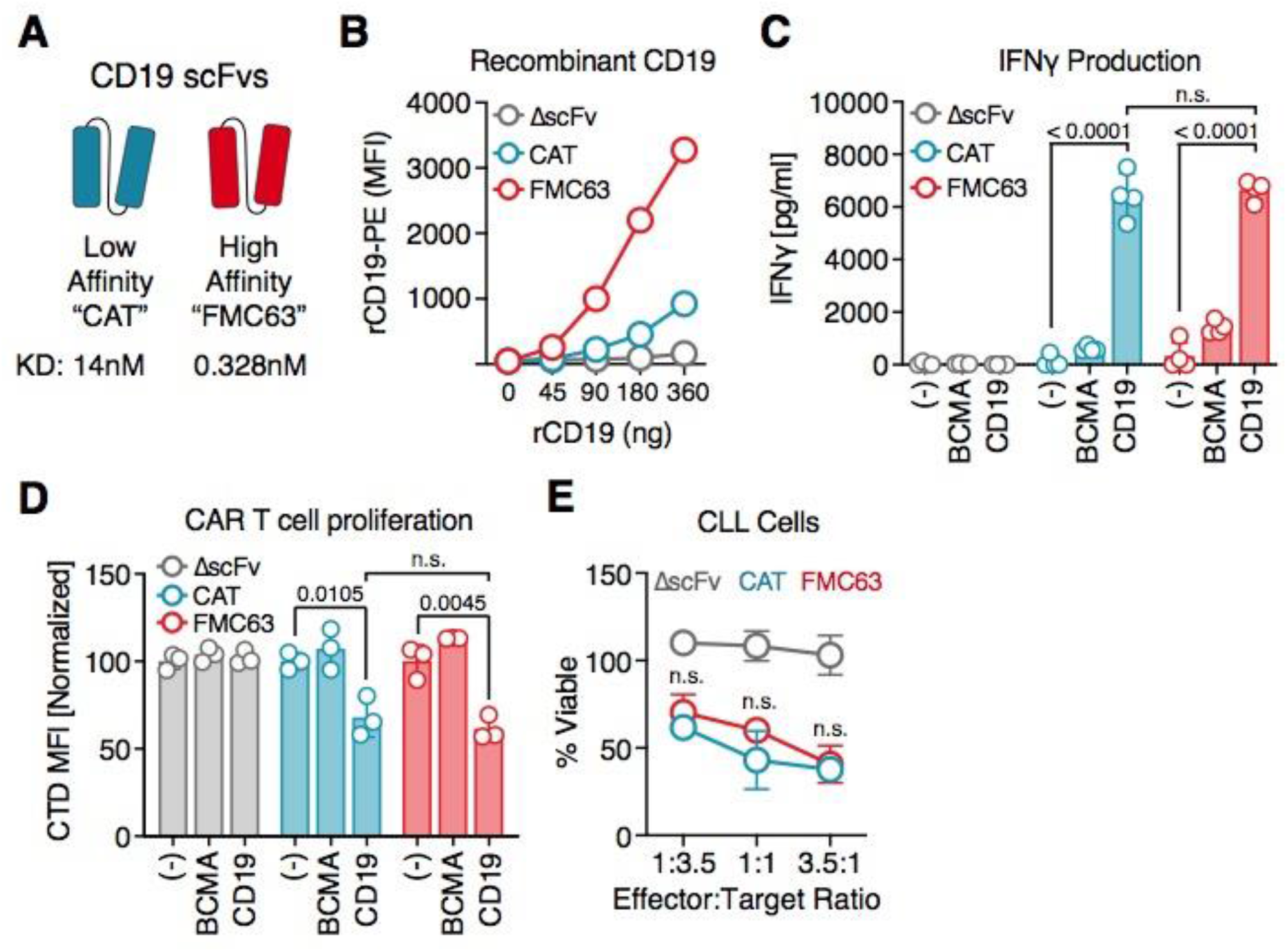
CAR affinity does not impact CAR T cell activation in the absence of trogocytosis. **(A)** Schema of high and low affinity CD19 scFvs. **(B)** CAR T cells were stained with increasing amounts of PE-labeled recombinant CD19 (Acro Biosystems # CD9-HP2H3) and assessed by flow cytometry. **(C)** CAR T cells were thawed and cultured in the presence of 40IU/mL IL2 before addition of CD19-coupled magnetic beads. After 48 hours, supernatants were harvested and IFN*γ* concentrations determined by ELISA. Data represent mean ± SD from four replicates. Statistical significance was determined by two-sided Student’s *t* test. **(D)** CAR T cells were cultured for 5 days without IL2 before staining with CellTrace Far Red dye and addition of CD19-coupled magnetic beads and 40IU/mL IL2. Dye dilution was determined after 5 days by flow cytometry. Data represent mean ± SD from three replicates. Statistical significance was determined by two-sided Student’s *t* test. **(E)** CAR T cells were cultured with primary CLL cells at varying effector:target ratios for 4 hours. CLL cell viability was assessed by flow cytometry using counting beads. Data represent mean ± SD from three replicates. Statistical significance was determined by two-sided Student’s *t* test.

### Low affinity CAR T cells show minimal trogocytosis across different B cell malignancies and target antigens

We next set out to determine the effect of reduced CAR affinity on trogocytosis. Using primary CLL cells from a relapsed patient, we observed that tumor cells cocultured with CAT CAR T cells had lost significantly less CD19 than those cocultured with FMC63 CAR T cells (Fig. 4A). Importantly, while treatment with FMC63 CAR T cells resulted in a substantial CD19-negative population, the remaining lymphoma cells treated with CAT CAR T cells still expressed intermediate to high levels of CD19, preventing the emergence of antigen-negative tumor cells (Fig. 4A). Importantly, we found that this effect is not specific to CLL cells, as primary DLBCL (Fig. 4B) and MCL (Fig. 4C) cells showed even less reduction in surface antigen expression when cocultured with low-affinity CAT CAR T cells compared to FMC63 CAR T cells. We next analyzed if reduced trogocytosis by low-affinity CAR T cells is specific to CD19. To this end, we generated single amino acid substitution high affinity (2D3.hi) and low affinity (2D3.lo) variants of our previously described CD229-specific scFv 2D3 (*31*). We then cocultured these variant CAR T cells with the CD229^+^ DLBCL cell line DB (Fig. 4D) (*31, 32*). We found that low-affinity CD229 CAR T cells stripped significantly less CD229 from DB cells when compared to high affinity CD229 CAR T cells, suggesting that the role of CAR affinity in CAR T cell-mediated trogocytosis is target independent (Fig. 4E).

**Figure 4.**
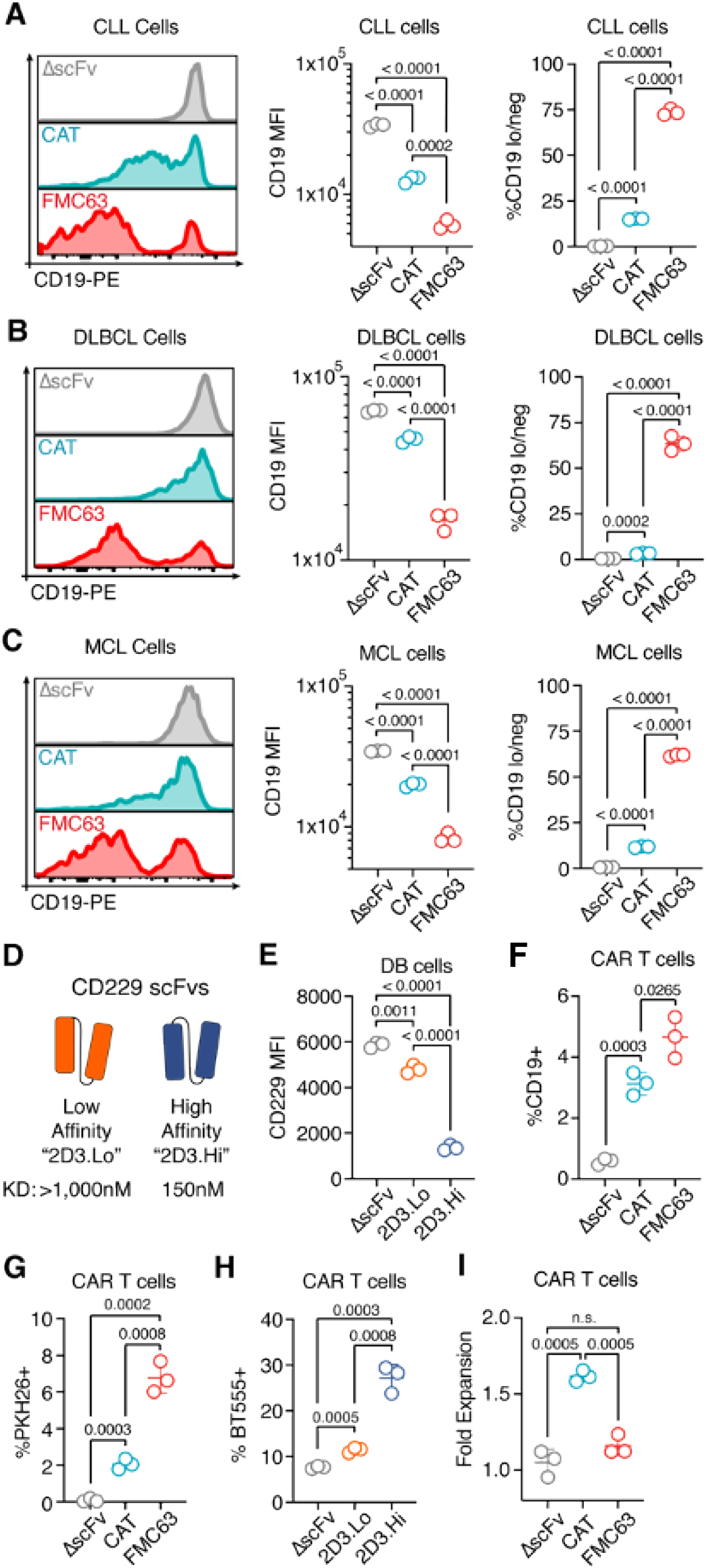
Low affinity CAR T cells exhibit reduced trogocytosis and fratricide. CD19 expression (left), CD19 MFI (middle) and % CD19 low/negative (right) **(A)** CLL, **(B)** DLBCL and **(C)** MCL cells following a 4 hour coculture with CAR T cells at an effector:target ratio of 4:1. **(D)** Schema of high and low affinity CD229 scFvs. **(E)** CD229 expression on DB cells labeled with CellTrace Far Red dye following a 3h coculture with CD229 CAR T cells at an effector:target ratio of 4:1. **(F)** % CD19+ CAR T cells following a 15 minute coculture with primary CLL cells at an effector:target ratio of 1:10. **(G)** Membrane transfer to CAR T cells following a 30 min coculture with PKH26 stained CLL cells at an effector:target ratio of 1:1. **(H)** Membrane transfer to CAR T cells following a 5hr coculture with BioTracker555 stained DB cells at an effector:target ratio of 4:1. All data including statistical analysis represent mean ± SD from three replicates. Statistical significance was determined by two-sided Student’s *t* test. **(I)** CAR T cell expansion following a 24 hour coculture with primary CLL cells at an effector:target ratio of 1:1 normalized to CAR T cell counts following a 24 hour coculture without CLL cells. CD19 expression (left), CD19 MFI (middle) and % CD19 low/negative (right) cells following a 4 hour coculture with CAR T cells at an effector:target ratio of 4:1.

### Reduced trogocytosis and increased persistence by low affinity CAR T cells

Finally, we analyzed the effect of affinity on trogocytosis-mediated CAR T cell persistence and fratricide. Co-culturing low-affinity CAT CAR T cells and high-affinity FMC63 CAR T cells with CD19^+^ CLL cells, we found that CAT CAR T cells had acquired substantially less CD19 (Fig. 4F) and less tumor cell membrane (Fig. 4G) than FMC63 CAR T cells (*29*). Similarly, low-affinity CD229 CAR T cells had also acquired substantially less tumor cell membrane than high-affinity CD229 CAR T cells (Fig. 4H). To determine if this reduction in trogocytosis also resulted in reduced fratricide of CAT CAR T cells, we determined CAR T cell numbers at the end of a 24h coculture with CLL cells. Indeed, we found that CAT CAR T cells, despite similar proliferative activity in response to CD19-coated beads, showed significantly greater expansion than FMC63 CAR T cells after coculture with CD19^+^ tumor cells (Fig. 4I), indicating that FMC63-based CAR T cells had exhibited significantly increased fratricide compare to CAT CAR T cells.

## DISCUSSION

All currently FDA-approved CD19 CAR T cell products rely on the high-affinity CD19 antibody FMC63 for antigen recognition. Using primary patient samples, we show that FMC63-based CAR T cells confer substantial antigen loss in all major tumor types currently approved or under clinical investigation for treatment with CD19 CAR T cells, potentially fueling a reservoir of persistent antigen-negative tumor cells. In addition, our observation of the rapid emergence and subsequent contraction of a CD19^+^ CAR T cell population, an increase in apoptosis in high-affinity CD19 CAR T cells, and the significantly increased number of low-affinity CAR T cells after tumor cell co-culture strongly indicate a substantial effect of trogocytosis-mediated fratricide on CAR T cell persistence. Our finding that trogocytosis is a very rapid and, as has been previously shown (*23*), transient process, likely also complicates its clinical validation in the absence of a more mechanistic understanding, e.g. allowing its targeted inhibition. However, our data also indicate that, at least in the case of CD19 CAR T cells, a clinically tested solution to the high levels of trogocytosis observed with FMC63-based CAR T cells is already available. We show that is possible to robustly sever trogocytosis and its deleterious effects from anti-tumor activity and other major T cell functions by using the low affinity CAT CAR construct (*33*). In addition to reduced trogocytosis, there may be other benefits to using low-affinity CAR binding domains, such as increased selectivity (*27, 34, 35*) and reduced exhaustion (*36*). We hypothesize that while the identification of low affinity constructs with equal anti-tumor activity may not be possible in all settings, e.g. due to low antigen density on tumor cells or selectivity requirements, the degree of trogocytosis resulting from differences in antibody affinity will likely become a key consideration in the development of novel CAR constructs.

## METHODS

### Cell lines and primary samples

NALM6 cells were purchased from DSMZ and cultured at 37°C/5% CO_2_ in ATCC-formulated RPMI-1640 (Gibco #A10491-01) supplemented with 10% FCS (Gibco #10437028) and 50U/mL penicillin/streptomycin. NALM6 cells were transduced with pHIV-Luc-ZsGreen lentivirus (Addgene #39196) and sorted by FACS prior to use. Primary patient samples were prepared as follows: whole blood or bone marrow aspirate was collected and mononuclear cells were isolated by density gradient using Ficoll-Paque (GE #17-1440-02), samples were cryopreserved until analysis.

### CAR constructs and virus production

CAR T cells were generated using lentiviral or gammaretroviral transduction. Second generation CAR constructs based on FMC63 (*37*), CAT (*33, 38*), or 2D3 (*31*) using the 4-1BB costimulatory and CD3*ζ* signaling domains, as well as the CD8*α* hinge and transmembrane domains were cloned into pRRLSIN.cPPT.PGK-GFP.WPRE (Addgene # 12252) or SFG.CNb30_opt.IRES.eGFP (Addgene # 22493) backbones. VSV-G-pseudotyped lentivirus was obtained by co-transfection of Lenti-X 293T cells (Takara) with pRRL-based transfer plasmids as well as pMD2.G (Adgene # 12259) and psPAX2 (Addgene # 12260) using Lipofectamine 2000 (Thermo) according to the manufacturer’s instructions. Amphotropic gammaretrovirus was generated by transfection of Phoenix-Ampho cells (ATCC # CRL-3213) with SFG-based transfer plasmids using Lipofectamine 2000 according to the manufacturer’s instructions. Virus-containing supernatants were concentrated with Lenti-X Concentrator or Retro-X concentrator (Takara), respectively, and 24 well plates coated with Retronectin (Takara # T100B) were incubated with concentrated viruses according to the manufacturer’s instructions.

### CAR T cell production

Buffy coats from healthy donors were obtained from the Blood Centers of America or the New York Blood Center and peripheral blood mononuclear cells (PBMC) were isolated by density gradient using Ficoll-Paque. T cells were cultured in AIM V (Thermo) supplemented with 5% human serum (Sigma #H3667). PBMCs were stimulated for 2 days with CD3/CD28 T cell activation beads (Thermo # 11131D) in the presence of 40IU/mL IL2 (R&D Systems # 202-IL-010) and incubated at 37°C/5% CO_2_. Bead-stimulated cells were transferred to Retronectin-coated virus-containing plates and incubated overnight. Transduction was repeated the next day before counting and diluting cells to 0.4×10^6^ cells/ml. After the second transduction cells were grown for an additional 7 days before removing beads using a DynaMag-15 magnet (Thermo). IL-2 was replenished every 2 days to 40IU/mL. Cells were frozen in 90% FCS/10% DMSO in liquid nitrogen until needed.

### *In vitro* trogocytosis assay

CAR T cells were cocultured with target cells at a defined effector:target ratio. For some experiments, target cells were first labelled with membrane dyes PKH26 (Sigma # MINI26) or BioTracker 555 (Sigma # SCT107) according to the manufacturer’s instructions. Cells were stained with the antibody panel described in Supplemental Table 1, 500ng/mL DAPI (Invitrogen # D1306) and counting beads (Thermo # PCB 100) were added following staining and cells were analyzed on a LSRFortessa or an LSR II flow cytometer (BD). Effector:target ratios and coculture times are listed in the figure legends. Gating schemes can be found in supplemental figure 3.

### Western blot

ΔscFv or CD19 CAR T cells were cocultured with primary CLL cells at an effector:target ratio of 4:1 for 4 hours. Tumor cells were sorted by FACS as CD3-GFP- and were lysed in RIPA buffer (Thermo #89900) containing protease inhibitor (Thermo # A32963). Protein concentration was determined by a bicinchoninic acid assay (Thermo # 23225) and lithium dodecyl sulfate buffer (Thermo # NP0007) was then used to denature the lysates. Equal amounts of total protein were loaded onto a 4-12% bis-tris gel and proteins were separated by sodium dodecyl sulfate– polyacrylamide gel electrophoresis and then transferred onto a nitrocellulose membrane.

Membranes were blocked with 5% non-fat milk-TBST for 1 hour at room temperature followed by incubation with anti-human CD19 antibody at 1:500 (CST # 90176S) or anti-human β-Actin clone 8H10D10 antibody at 1:750 (CST # 3700S) overnight at 4°C. Membranes were washed and then incubated in HRP-linked, anti-rabbit (CST #7074S) or anti-mouse (CST #7076S) IgG antibody at 1:10,000 for 1 hour at room temperature. Membranes were washed and proteins were visualized by enhanced chemiluminescence (Thermo Pierce ECL Plus # 32132) and captured on X-ray film (Thermo # 34090).

### *In vivo* trogocytosis model

7-8-week-old female NSG mice were randomly grouped and sublethally irradiated at 5cGy on the morning of day −2 followed by i.v. administration of 2.5×10^5^ NALM6-Luc cells. Mice were administered 3.3 mg D-Luciferin in the morning of day 3 and luminescence was recorded after 10 minutes by an In Vivo Imaging System (IVIS, Perkin Elmer) followed by i.v. injection of 2.5×10^6^ ΔscFv or FMC63 CAR T cells. All injections and monitoring by IVIS were performed by the preclinical research resource core at the University of Utah, a blinded third party. Tumor progression was monitored by IVIS on the morning of day 10 and cervical lymph nodes were collected in the afternoon. Lymph nodes were stained for CD19 expression using the panel described in supplemental table 1 following incubation with FcR blocking reagent (Miltenyi Biotec # 130-092-575). Cells were analyzed on a BD LSRFortessa. Gating schemes can be found in supplemental figure 3.

### CD19 bead stimulation assays

For CAR T cell proliferation assays, ΔscFv, FMC63, and CAT CAR T cells were stained with CellTrace Far Red dye (Thermo # C34564) according to the manufacturer’s instructions. CAR T cells were starved of IL-2 for 5 days before the addition of CD19-coated (Acro Biosystems # MBS-K005) or BCMA-coated (Acro Biosystems # MBS-K004) paramagnetic beads. 40IU/mL IL-2 was added to cultures together with beads, and replenished every 2 days. Dye dilution was determined after 5 days on an LSR II flow cytometer (BD). IFN*γ* concentrations in cell culture supernatants were determined after 48h by ELISA (Biolegend # 430104) according to the manufacturer’s instructions. Flow cytometry data was analyzed using FlowJo 10 (BD).

### Study approval

Informed consent was obtained from all patients and samples were collected with the approval of the Institutional Review Boards of the University of Utah (IRB #45880) and Tisch Cancer Institute (IRB #13-1347, 17-2164, 17-0554). All animal procedures were conducted under protocol #16-05007, approved by the Institutional Animal Care and Use Committee at the University of Utah.

### Statistical analyses

Significance of differences in cell numbers, cytokine levels, and mean fluorescence intensity were calculated by two-tailed Student’s t-test. All statistical tests were performed using Prism 9 (GraphPad Software). Results were considered significant when p or adjusted p < 0.05.

### Supplemental material

There are 3 supplementary figures and 1 supplementary table included here. Supplementary figure 1 describes *in vitro* trogocytosis by CD19 CAR T cells; supplementary figure 2 describes the production of the FMC63 and CAT CAR T cells; supplementary figure 3 describes gating schemes for flow cytometry analysis; supplementary table 1 describes antibodies used for flow cytometry.

## AUTHOR CONTRIBUTIONS

MLO conceived the project, performed experiments, analyzed data and wrote the manuscript. EVRM generated the 2D3 affinity variants and analyzed data. SVR and APR analyzed data. ALW and DA wrote the manuscript and analyzed data. JDB provided CLL, DLBCL and MCL patient samples. TL conceived the project, generated CAR T cells, performed the CD19 bead stimulation experiments, performed the CD229 CAR T cell trogocytosis experiments, analyzed data and wrote the manuscript.

## ACKNOWLEDGEMENTS

We thank the preclinical research resource at the Huntsman Cancer Institute for handling and treating the animals for our mouse model, the flow cytometry core for assistance in cell sorting, and the Huntsman Cancer Institute tissue banking core for providing CLL patient samples. pHIV-Luc-ZsGreen was a gift from Bryan Welm (Addgene #39196), pRRLSIN was a gift from Didier Trono (Addgene # 12252), and SFG.CNb30_opt.IRES.eGFP was a gift from Martin Pule (Addgene # 22493).

## REFERENCES

1. J. Pan et al., Frequent occurrence of CD19-negative relapse after CD19 CAR T and consolidation therapy in 14 TP53-mutated r/r B-ALL children. Leukemia 34, 3382–3387 (2020).

2. S. L. Maude et al., Chimeric antigen receptor T cells for sustained remissions in leukemia. The New England journal of medicine 371, 1507–1517 (2014).

3. J. H. Park et al., Long-Term Follow-up of CD19 CAR Therapy in Acute Lymphoblastic Leukemia. N Engl J Med 378, 449–459 (2018).

4. K. Wudhikarn et al., Interventions and outcomes of adult patients with B-ALL progressing after CD19 chimeric antigen receptor T-cell therapy. Blood 138, 531–543 (2021).

5. V. Plaks et al., CD19 target evasion as a mechanism of relapse in large B-cell lymphoma treated with axicabtagene ciloleucel. Blood, (2021).

6. K. Sakamoto et al., Negative CD19 expression is associated with inferior relapse-free survival in children with RUNX1-RUNX1T1-positive acute myeloid leukaemia: results from the Japanese Paediatric Leukaemia/Lymphoma Study Group AML-05 study. Br J Haematol 187, 372–376 (2019).

7. A. Bukhari et al., Rapid relapse of large B-cell lymphoma after CD19 directed CAR-T-cell therapy due to CD-19 antigen loss. Am J Hematol 94, E273–E275 (2019).

8. T. Rabilloud et al., Single-cell profiling identifies pre-existing CD19-negative subclones in a B-ALL patient with CD19-negative relapse after CAR-T therapy. Nat Commun 12, 865 (2021).

9. J. Cao et al., Potent anti-leukemia activities of humanized CD19-targeted Chimeric antigen receptor T (CAR-T) cells in patients with relapsed/refractory acute lymphoblastic leukemia. Am J Hematol 93, 851–858 (2018).

10. E. J. Orlando et al., Genetic mechanisms of target antigen loss in CAR19 therapy of acute lymphoblastic leukemia. Nat Med 24, 1504–1506 (2018).

11. J. Fischer et al., CD19 Isoforms Enabling Resistance to CART-19 Immunotherapy Are Expressed in B-ALL Patients at Initial Diagnosis. J Immunother 40, 187–195 (2017).

12. E. Sotillo et al., Convergence of Acquired Mutations and Alternative Splicing of CD19 Enables Resistance to CART-19 Immunotherapy. Cancer Discov 5, 1282–1295 (2015).

13. E. Jacoby et al., CD19 CAR immune pressure induces B-precursor acute lymphoblastic leukaemia lineage switch exposing inherent leukaemic plasticity. Nat Commun 7, 12320 (2016).

14. J. Y. Huang, Y; Sepulveda, WS; Hwang, I; Peterson, P; Jackson, MR; Sprent, J; Cai, Z, TCR-Mediated Internalization of Peptide-MHC Complexes Acquired by T Cells. Science 286, (1999).

15. R. Brown et al., CD86+ or HLA-G+ can be transferred via trogocytosis from myeloma cells to T cells and are associated with poor prognosis. Blood 120, 2055–2063 (2012).

16. J. LeMaoult et al., Immune regulation by pretenders: cell-to-cell transfers of HLA-G make effector T cells act as regulatory cells. Blood 109, 2040–2048 (2007).

17. A. Aucher, E. Magdeleine, E. Joly, D. Hudrisier, Capture of plasma membrane fragments from target cells by trogocytosis requires signaling in T cells but not in B cells. Blood 111, 5621–5628 (2008).

18. M. S. Ford McIntyre, K. J. Young, J. Gao, B. Joe, L. Zhang, Cutting edge: in vivo trogocytosis as a mechanism of double negative regulatory T cell-mediated antigen-specific suppression. J Immunol 181, 2271–2275 (2008).

19. E. P. Dopfer, S. Minguet, W. W. Schamel, A new vampire saga: the molecular mechanism of T cell trogocytosis. Immunity 35, 151–153 (2011).

20. S. S. Somanchi, A. Somanchi, L. J. Cooper, D. A. Lee, Engineering lymph node homing of ex vivo-expanded human natural killer cells via trogocytosis of the chemokine receptor CCR7. Blood 119, 5164–5172 (2012).

21. R. Valgardsdottir et al., Human neutrophils mediate trogocytosis rather than phagocytosis of CLL B cells opsonized with anti-CD20 antibodies. Blood 129, 2636–2644 (2017).

22. M. Kawashima et al., PD-L1/L2 protein levels rapidly increase on monocytes via trogocytosis from tumor cells in classical Hodgkin lymphoma. Leukemia 34, 2405–2417 (2020).

23. M. Hamieh et al., CAR T cell trogocytosis and cooperative killing regulate tumour antigen escape. Nature 568, 112–116 (2019).

24. S. I. Richardson et al., IgG3 enhances neutralization potency and Fc effector function of an HIV V2-specific broadly neutralizing antibody. PLoS Pathog 15, e1008064 (2019).

25. B. Chung et al., Antigen-specific inhibition of high-avidity T cell target lysis by low-avidity T cells via trogocytosis. Cell Rep 8, 871–882 (2014).

26. M. Chmielewski, A. Hombach, C. Heuser, G. P. Adams, H. Abken, T Cell Activation by Antibody-Like Immunoreceptors: Increase in Affinity of the Single-Chain Fragment Domain above Threshold Does Not Increase T Cell Activation against Antigen-Positive Target Cells but Decreases Selectivity. The Journal of Immunology 173, 7647 (2004).

27. X. Liu et al., Affinity-Tuned ErbB2 or EGFR Chimeric Antigen Receptor T Cells Exhibit an Increased Therapeutic Index against Tumors in Mice. Cancer Res 75, 3596–3607 (2015).

28. F. Turatti et al., Redirected activity of human antitumor chimeric immune receptors is governed by antigen and receptor expression levels and affinity of interaction. J Immunother 30, 684–693 (2007).

29. S. Ghorashian et al., Enhanced CAR T cell expansion and prolonged persistence in pediatric patients with ALL treated with a low-affinity CD19 CAR. Nat Med 25, 1408–1414 (2019).

30. C. Roddie et al., Durable Responses and Low Toxicity After Fast Off-Rate CD19 Chimeric Antigen Receptor-T Therapy in Adults With Relapsed or Refractory B-Cell Acute Lymphoblastic Leukemia. J Clin Oncol, JCO2100917 (2021).

31. S. V. Radhakrishnan et al., CD229 CAR T cells eliminate multiple myeloma and tumor propagating cells without fratricide. Nat Commun 11, 798 (2020).

32. E. Vander Mause et al., Highly selective CD229 chimeric antigen receptor generated using affinity tuning platform. Submitted.

33. S. Ghorashian et al., Enhanced CAR T cell expansion and prolonged persistence in pediatric patients with ALL treated with a low-affinity CD19 CAR. Nature Medicine 25, 1408–1414 (2019).

34. S. Arcangeli et al., Balance of Anti-CD123 Chimeric Antigen Receptor Binding Affinity and Density for the Targeting of Acute Myeloid Leukemia. Molecular Therapy 25, 1933–1945 (2017).

35. S. Park et al., Micromolar affinity CAR T cells to ICAM-1 achieves rapid tumor elimination while avoiding systemic toxicity. Scientific Reports 7, 14366 (2017).

36. A. P. Singh et al., Development of a quantitative relationship between CAR-affinity, antigen abundance, tumor cell depletion and CAR-T cell expansion using a multiscale systems PK-PD model. MAbs 12, 1688616 (2020).

37. I. C. Nicholson et al., Construction and characterisation of a functional CD19 specific single chain Fv fragment for immunotherapy of B lineage leukaemia and lymphoma. Mol Immunol 34, 1157–1165 (1997).

38. C.G. Amrolia P, Ghorashian S, Kramer A, Mekkaoui L, Pule M, USPTO, Ed. (US, 2019).

