## Supplementary Material for "Low-affinity CAR T cells exhibit reduced trogocytosis, preventing fratricide and antigen-negative tumor escape while preserving anti-tumor activity"

**SUPPLEMENTAL DATA**

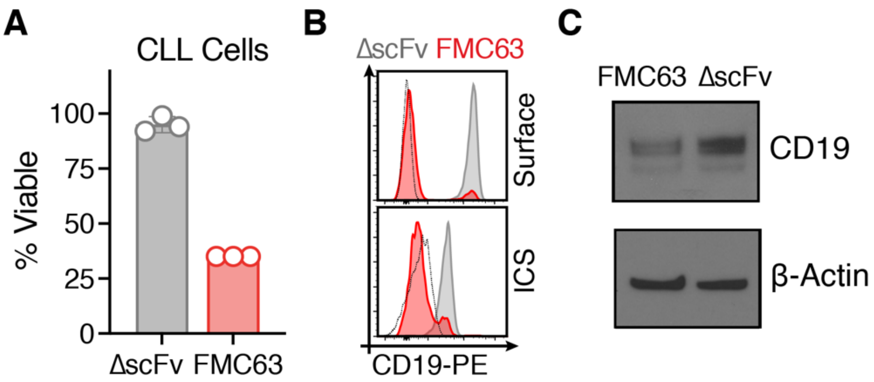


**Supplementary Figure 1: CD19 CAR T cell-mediated trogocytosis.** ΔscFv (grey) and FMC63 (red) CAR T cells were cocultured with primary CLL cells for 4 hours at an effector:target ratio of 4:1. **(A)** Viability of CLL cells following coculture as determined by flow cytometry. Data represent technical replicates (N=3). **(B)** Surface (top) and intracellular (bottom) staining of CD19 on CLL cells following coculture as determined by flow cytometry. Dotted line denotes isotype staining. **(C)** Expression of total CD19 and β-Actin in primary CLL cells after co-culture with CD19 CAR T cells or ΔscFv CAR T cells as determined by western blot.

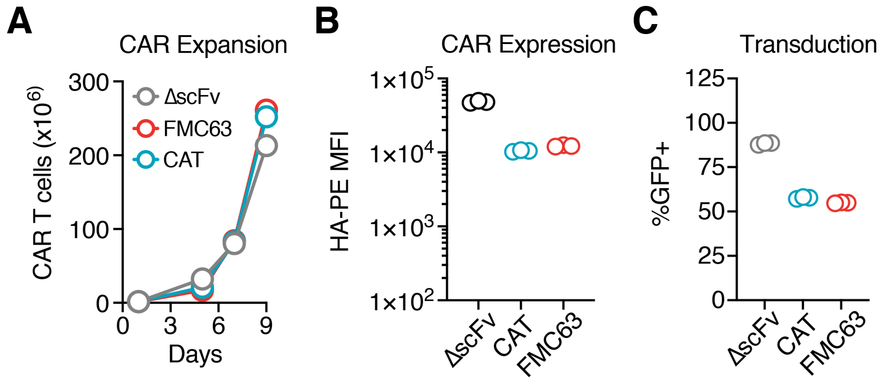


**Supplementary Figure 2: Characterization of low and high affinity CD19 CAR T cells.** High (FMC63) and Low (CAT) affinity CD19 CAR T cells as well as CAR T cells without a binding domain (ΔscFv) were generated using gammaretroviral transduction. **(A)** CAR T cells were expanded for 9 days and supplemented with 40IU/ml IL2 every two days before freezing. CAR T cell numbers were determined after trypan blue staining on a Countess II cell counter. Plot shows a representative result from 3 independent productions. **(B)** CAR surface expression was determined after staining with an anti-HA-PE antibody on an LSR II flow cytometer. Data represent technical replicates (N=3). **(C)** CAR T cell transduction rate was determined by GFP reporter expression using an LSR II flow cytometer. Data represent technical replicates (N=3).

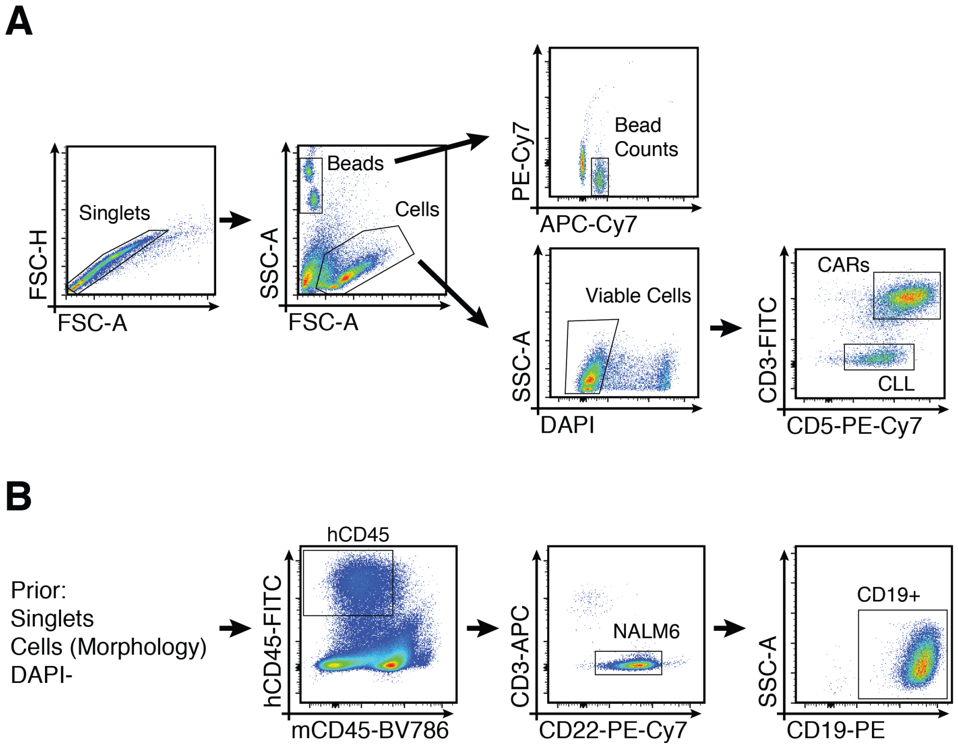


**Supplementary Figure 3: Flow cytometry gating schemes. (A)** Gating scheme for *in vitro* cytotoxicity and trogocytosis assays. **(B)** Gating scheme for *in vivo* trogocytosis assay, cells are pre-gated on singlets, cell morphology and DAPI- as shown in A.

| Experiment | Target | Fluorophore | Manufacturer | Catolog # |
|---|---|---|---|---|
| *In vivo* trogocytosis model (Fig 1D) | CD3 | APC | Biolegend | 300412 |
| | CD19 | PE | Biolegend | 302208 |
| | CD22 | PE-Cy7 | Biolegend | 302514 |
| | huCD45 | FITC | Biolegend | 304006 |
| | mseCD45 | BV-786 | Biolegend | 110743 |
| Trogocytosis assays (Fig 1E-F, Fig 2E-G, I-K) | CD3 | APC | Biolegend | 300412 |
| | CD19 | PE | Biolegend | 302208 |
| | CD5 | PE-Cy7 | Biolegend | 300622 |
| | CD5 | FITC | Biolegend | 302303 |
| CAR T cell annexinV stain (Fig 1G) | AnnexinV | Pacific Blue | Biolegend | 640926 |
| | CD3 | APC | Biolegend | 300412 |
| | CD19 | PE | Biolegend | 302208 |
| Trogocytosis primary sample panel (Fig 1I) | CD3 | APC | Biolegend | 300412 |
| | CD5 | BV421 | Biolegend | 300626 |
| | CD10 | BV421 | Biolegend | 312218 |
| | CD19 | PE | Biolegend | 302208 |
| | CD20 | FITC | Biolegend | 302303 |
| rCD19 stain (Fig 2B) | PE labeled recombinant CD19 | | Acro Biosystems | CD9-HP2H3 |
| HA Stain | HA | PE | Biolegend | 901517 |
| CD229 trogo (Fig 2M) | CD229 | APC | Invitrogen | 17-2299-42 |

**Supplementary Table 1: Antibodies used for flow cytometry**
